# Uncovering Lasonolide A biosynthesis using genome-resolved metagenomics

**DOI:** 10.1101/2022.05.23.493085

**Authors:** Siddharth Uppal, Jackie L. Metz, René K.M. Xavier, Keshav Nepal, Dongbo Xu, Guojun Wang, Jason C. Kwan

## Abstract

Invertebrates, in particular sponges, have been a dominant source of new marine natural products. For example, lasonolide A (LSA) is a potential anti-cancer molecule isolated from the marine sponge *Forcepia* sp., with nanomolar growth inhibitory activity and a unique cytotoxicity profile against the National Cancer Institute 60 cell line screen. Here, we identified the putative biosynthetic pathway for LSA. Genomic binning of the *Forcepia* sponge metagenome revealed a gram-negative bacterium belonging to the phylum Verrucomicrobia as the candidate producer of LSA. Phylogenetic analysis showed this bacterium, herein named *Candidatus* Thermopylae lasonolidus, only has 88.78% 16S rRNA identity with the closest relative *Pedosphaera parvula* Ellin514, indicating it represents a new genus. The lasonolide A (*las*) biosynthetic gene cluster (BGC) was identified as a *trans-*AT polyketide synthase (PKS) pathway. When compared with its host genome, the *las* BGC exhibits a significantly different GC content and penta-nucleotide frequency, suggesting a potential horizontal acquisition of the gene cluster. Furthermore, three copies of the putative *las* pathway were identified in the candidate producer genome. Differences between the three *las* repeats were observed including the presence of three insertions, two single-nucleotide polymorphisms and the absence of a stand-alone acyl carrier protein in one of the repeats. Even though the Verrucomicrobial producer shows signs of genome-reduction, its genome size is still fairly large (about 5Mbp) and when compared to its closest free-living relative contains most of the primary metabolic pathways, suggesting that it is in the early stages of reduction.

**Importance:** While sponges are valuable sources of bioactive natural products, a majority of these compounds are produced in small amounts by uncultured symbionts, hampering the study and clinical development of these unique compounds. Lasonolide A (LSA), isolated from marine sponge *Forcepia* sp., is a cytotoxic molecule active at nanomolar concentrations and causes premature chromosome condensation, blebbing, cell contraction and loss of cell adhesion, indicating a novel mechanism of action and making it a potential anti-cancer drug lead. However, its limited supply hampers progression to clinical trials. We investigated the microbiome of *Forcepia* sp. using culture-independent DNA sequencing to uncover how an uncultured bacterium produces LSA. This provides future opportunities for heterologous expression and cultivation efforts that may minimize LSA’s supply problem.

## Introduction

Lasonolide A (LSA) is a cytotoxic polyketide derived from the marine sponge *Forcepia* sp. (**Fig. 1A and 1B**) (1). Out of its analogs (B–G) (**Fig. 1C**), LSA is the most potent (2) and exhibits IC_50_ values in the nanomolar range against certain cell lines in the National Cancer Institute 60 cell line screen (3). Furthermore, its unique mechanism of action - induction of premature chromosome condensation, loss of cell adhesion, activation of the RAF1 kinase in Ras pathway, along with cell blebbing and contraction (3–5) - makes it a promising candidate as a scaffold for future pharmaceutical development. However, a major challenge to its clinical development is the lack of availability. Scarcity and limited accessibility of the sponge prevent it from being a sustainable source of lasonolide A. Furthermore, the chemical synthesis of LSA is tedious and has poor yields, limiting its scalability (6–8).

**Fig. 1.**
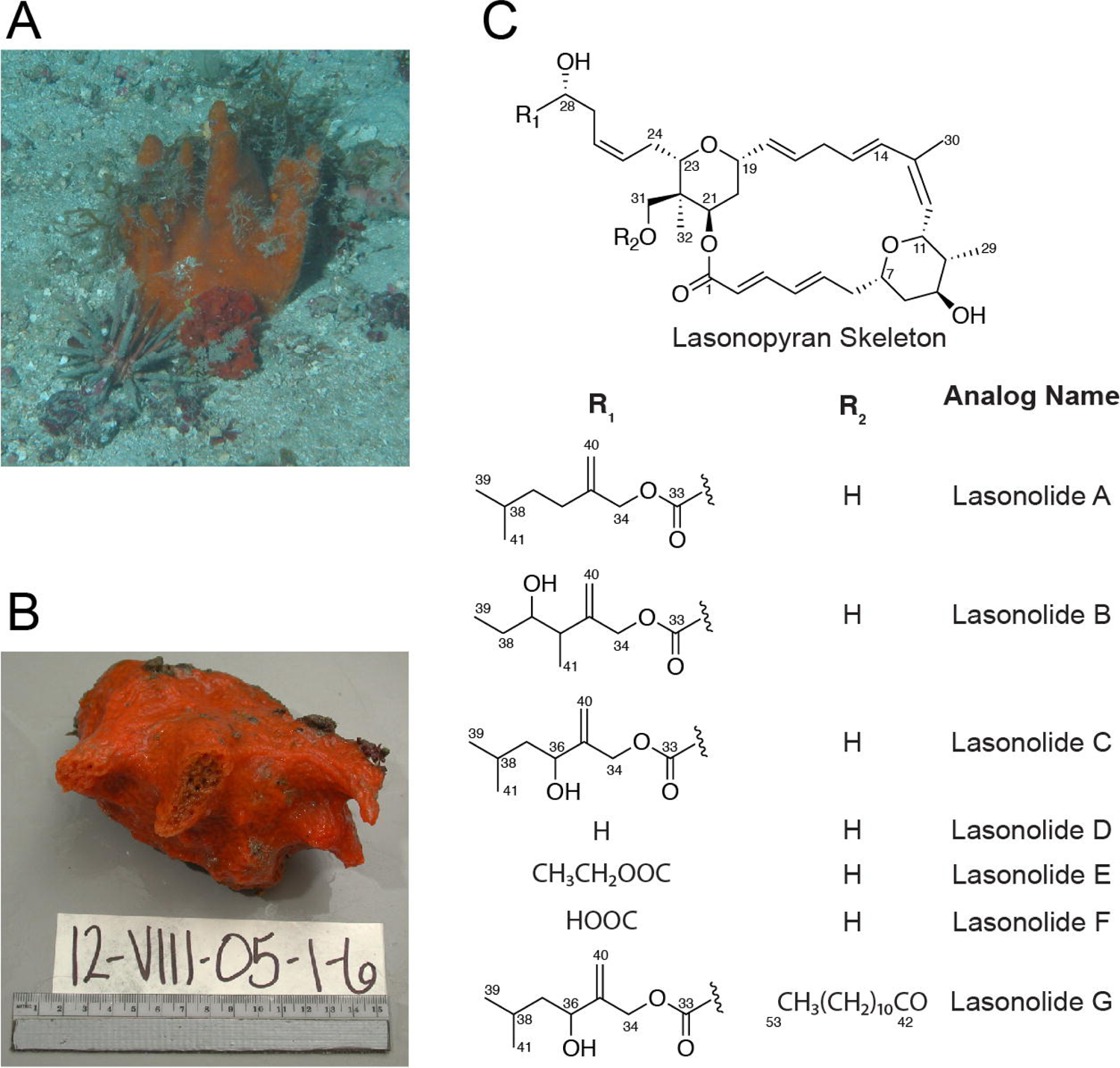
(A) Sponge *Forcepia* sp. as seen in the field. (B) The *Forcepia* sp. specimen used for DNA extraction (sample ID: 12-VIII-05-1-6). Photo credit: HBOI Marine Biomedical and Biotechnology Program. (C) The chemical structures of lasonolide A (LSA) and its analogs.

It is well known that bacteria living in a symbiotic relationship with higher animals are valuable sources of novel bioactive secondary metabolites (9). In many instances, these molecules serve a protective function for the host but the identity of the microbial producer remains unknown (9–12). Furthermore, attempts to isolate these associated microbes are hampered by low cultivation success; it is estimated less than 1% of bacteria are currently culturable from the environment (13–15). These drawbacks have created the need to genetically engineer surrogate hosts for the sustainable and sufficient production of the desired natural products in the laboratory. The first step in engineering microbes for production of bioactive compounds is to identify the genes responsible for natural product synthesis, which can be elucidated through metagenomic analysis and cloning (16, 17). Based on its potent antitumor activity, it is likely that LSA also acts as a chemical defense within its host sponge. The structure of LSA very likely arises from an assembly-line type polyketide synthase (PKS), rather than the iterative PKSs that predominate in fungi and other eukaryotes, and therefore the source is likely bacterial (18–20). Identifying the bacterium responsible for synthesizing LSA and elucidating its biosynthetic pathway will allow us to explore routes for LSA’s heterologous expression, and potentially facilitate the synthesis of analogs.

Here, we describe a *trans-*AT PKS pathway (*las*) that is likely responsible for the biosynthesis of LSA. Furthermore, the entire *las* BGC has been captured on five overlapping fosmid clones and reassembled for the purpose of heterologous expression. We propose that the *las* BGC is present in a yet uncultivated bacterium belonging to a novel genus under the phylum Verrucomicrobia. Additionally, evidence suggests the *las* BGC is repeated thrice within the Verrucomicrobia symbiont with minor sequence variations between them. We also suggest that the *las* BGC has been horizontally acquired and has a codon adaptation index comparable to that of highly expressed genes. Finally, we show that the Verrucomicrobia symbiont is in very early stages of genome reduction and is likely to further reduce its size.

## Results and discussion

### Identification and capture of *las* BGC

In our initial studies, we constructed a high-capacity metagenomic DNA library consisting of ∼600,000 *cfu* from *Forcepia* sp. sponges collected from the Gulf of Mexico (**Fig. 2A**) to search for potential *las* biosynthetic genes. The structure of LSA contains two tetrahydropyran rings and β-methylations (21, 22) at C-13 and C-35 (**Fig. 2B**). These structural features have been identified in a variety of *trans-*AT PKS pathways but are rarely found in *cis-*AT PKS systems (23, 24) thus hinting that LSA is produced by a *trans-*AT PKS pathway (24). Therefore, clade-guided degenerate primers targeted to conserved *trans*-AT PKS genes such as 3-hydroxy-3-methyglutaryl-CoA (HMG-CoA) synthase, free-standing ketosynthase (KS), acyl carrier protein (ACP), and two enoyl-CoA hydratases (ECH) were utilized for initial screening of the *Forcepia* fosmid library (**Table S1A**). From the metagenomic library, five fosmids were identified using these primers (fosmids 5-16, 6-71, 3-46, 1-80, and 4-77) resulting in the capture of approximately 48kb of the putative *las* BGC at its 3′ end (**Fig. S1A**). However, minimal progress was made toward capturing the remaining half of the BGC as primer walking failed to produce new hits in the region upstream of fosmid 5-16. Therefore, we sequenced the metagenome of *Forcepia* sp. and searched for *trans-*AT PKS BGCs. DNA was extracted from two different regions (referred to as Forcepia_v1 and Forcepia_v2) of the same sponge, and subjected to whole genome shotgun metagenomic sequencing. The reads were trimmed, assembled and then binned into metagenome assembled genomes (MAGs). The metagenomes were found to be abundant in Acidobacteria, Proteobacteria and Chloroflexota (**Fig. 2C and S1B**), with 56 and 55 MAGs recovered from the two metagenomes. Based on MiMAG (25) standards for completeness and contamination, 11 and 6 MAGs were high quality and 21 and 19 MAGs were medium quality, for Forcepia_v1 and Forcepia_v2, respectively (**Table S2**).

**Fig. 2.**
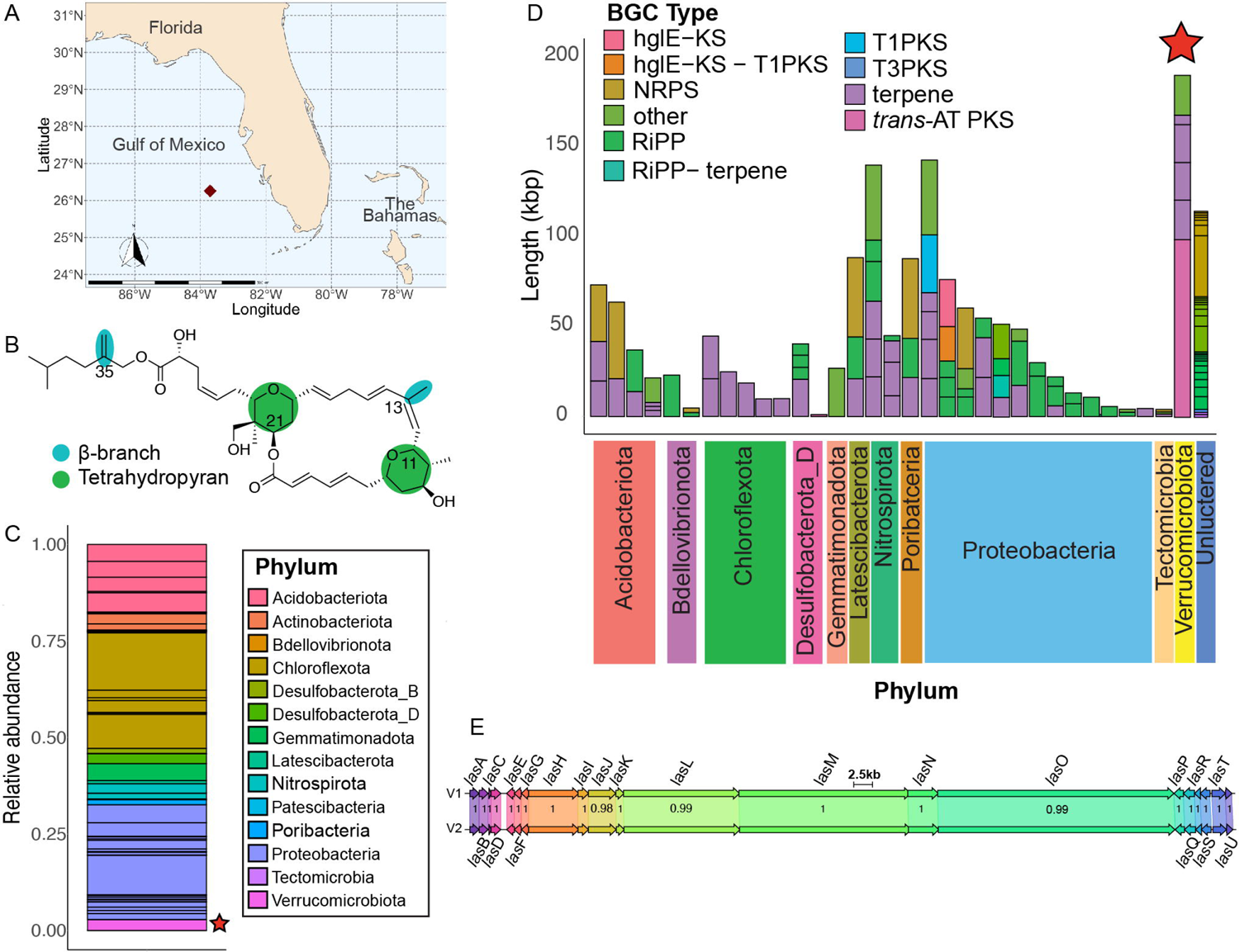
**(A)** Collection site of *Forcepia* sp. sponge (dark red diamond, 26.256573N, 83.702772W). **(B)** Features in lasonolide A (LSA) characteristic of biosynthesis by a *trans-*AT PKS pathway. **(C)** Relative abundance of different phyla in the sequenced Forcepia_v1 metagenome. Each block shows the relative abundance of each metagenome-assembled genome (MAG), with colors representing the phylum they belong to. The *las* biosynthetic gene cluster (BGC)-carrying bin is marked with a star. **(D)** BGC distribution in *Forcepia*_v1 sp. metagenome. AntiSMASH (27) annotations of bacterial contigs greater than 500 bp are shown. Each bar indicates a MAG. Bars have been grouped by phylum. The star represents the MAG possessing *las* BGC. BGC annotations have been simplified into polyketide synthase (PKS), Type 1 PKS, Type 3 PKS, *trans-*AT PKS, nonribosomal peptide synthetase (NRPS), ribosomally synthesized, post-translationally modified peptide (RiPP), hgIE-KS, hgIE-KS-T1PKS, terpenes, RiPP-terpene and others. **(E)** Comparison of *las* BGC_v1 and *las* BGC_v2 using clinker (28). V1 refers to *las* BGC_v1 while V2 refers to *las* BGC_v2. Numbers in the boxes indicate amino acid identity as a fraction of 1.

A tBLASTN (26) search of KS domains from publicly available *trans-*AT PKS pathways against our assembled metagenome was performed. In the case of Forcepia_v1, the top hits were all to a contig of length 98kbp labeled gnl|UoN|bin5_1_edit_8, thus strongly suggesting that this contig contains *trans-*AT PKS genes and may possess the potential LSA pathway. Contig gnl|UoN|bin5_1_edit_8 was manually inspected and corrected for sequence gaps (**Text S1**). With the exception of a 1.1kbp contig annotated as containing a *trans*-AT PKS pathway with a truncated condensation domain (in bin3674_131), analysis of the metagenome using AntiSMASH (27) (**Fig. 2D**) did not reveal any other BGC with plausible size and genes for the synthesis of LSA. Contig gnl|UoN|bin5_1_edit_128 (3.6 kbp) which was connected to the 5 end of gnl|UoN|bin5_1_edit_8 was found to contain a stand-alone ACP domain and about 47 amino acid residues which completed the terminal KS domain of gnl|UoN|bin5_1_edit_8 (see multiple repeats of the *las* BGC). Both the contigs were assembled together and annotation of genes and biosynthetic domains within this assembly re-affirmed that they are likely involved in LSA synthesis. We termed the gene cluster deemed relevant to LSA biosynthesis as *las* BGC_v1. Furthermore, the sequence of *las* BGC_v1 was also in alignment with fosmids identified from the metagenomic library. A screening strategy was then developed for isolating the previously missing 5′ end of the pathway from the metagenomic library (fosmids 5-41, 2-18 and 2-13) (**Fig. S1A and Table S1B**).

Inspection of the MAGs revealed that gnl|UoN|bin5_1_edit_8 binned with genome bin75_1. However, to our surprise, visual inspection of the assembly graph (**Fig. S1C**) in BANDAGE (29) indicated that gnl|UoN|bin5_1_edit_8 is present between contigs belonging to bin5_1 (phylum Verrucomicrobia). Furthermore, mapping the paired-end reads on the genomic bin (**Fig. S1D**) showed that multiple read pairs aligned across the contig junction. The terminal connections between contig gnl|UoN|bin5_1_edit_8 and several contigs in bin5_1 were verified via PCR (**Table S1C**) and Sanger sequencing of the amplicons using metagenomic DNA as the template. Based on this evidence gnl|UoN|bin5_1_edit_8 was manually placed with bin5_1, as well as additional contigs (**Text S1**).

In the case of Forcepia_v2, tBLASTN of KS domains hit to eight different contigs which could be assembled together (https://www.geneious.com) (**Fig. S1E**). Except for contig gnl|UoN|bin4_1_edit_10 the other seven contigs assembled into a single large contig of 102kbp (*las* BGC_v2). Similar to *las* BGC_v1, inspection of the assembly graph (**Fig. S1F)** and mapping of paired-end reads (**Fig. S1G**) revealed that contigs forming *las* BGC_v2 have been binned incorrectly and should be part of the bin4_1 (phylum Verrucomicrobia). As a result, the contigs comprising *las* BGC_v2, as well as additional contigs (**Text S1**) were manually placed with bin4_1. No other contig containing a *trans*-AT PKS pathway was identified in the metagenome (**Fig. S1H**).

Alignment of *las* BGC from both Forcepia_v1 and Forcepia_v2 using clinker (28) revealed that these pathways are highly similar (**Fig. 2E**). The amino acid identity is 100% for most of the genes except for *lasJLO* where the amino acid identity is 98.37%, 99.84% and 99.83% respectively. The slightly lower identity of *lasJLO* is due to the insertion sequence present in *las* BGC_v2 but absent in *las* BGC_v1. These insertion variants were later identified to be present in some repeats of *las* BGC_v1 as well.

The putative symbiont genome carrying the *las* BGC (Forcepia_v1 bin5_1 and Forcepia_v2 bin4_1) was identified to belong to phylum Verrucomicrobiota, order Pedosphaerales, and genus UBA2970 by GTDB-TK v1.5.0 (database r202) (30). Excluding the *las* genes, the ANI of Forcepia_v1 bin5_1 and Forcepia_v2 bin4_1 is 99.9%, suggesting little strain heterogeneity between the sites in the sponge, beyond a small amount perhaps attributable to sequencing errors. To our knowledge, this is the first time a *trans*-AT PKS BGC has been reported in an organism belonging to order Pedosphaerales. A phylogenetic tree of 51 different Verrucomicrobia genomes (**Fig. S1I**) placed the LSA producer in subdivision 3 (NCBI taxonomy). The closest relative of the symbiont with a publicly available genome is *Pedosphaera parvula* Ellin514 (GCA_000172555.1), with 88.78% identity to the 16S ribosomal RNA sequence. As per the 16S taxonomic cutoffs proposed by *Yarza* et al. (31), this represents a new genus within the family AAA164-E04 (as classified by GTDB-Tk (30)). We named the bacterium “*Candidatus* Thermopylae lasonolidus”: Thermopylae is a tribute to the 300 Spartan hoplites and other Greek soldiers that fought at the battle of Thermopylae. The Spartans fought to protect Greece from Persians and the LSA-producing bacterium with its three copies of the *las* BGC (see below) is proposed to be protecting the host sponge from predators. Lasonolidus suggests the bacterium is associated with lasonolide A and also rhymes with the Spartan king of the 300 hoplites, Leonidas. Despite being the putative producer of LSA, “*Ca* T. lasonolidus” is not highly abundant in the metagenome, having a relative abundance of just over 2.65% in Forcepia_v1 and 1.78% in Forcepia_v2 (**Fig. 2C and Fig. S1B**).

With the aid of metagenome sequence, additional fosmids covering the 5-end were acquired (Fig. S1), which enabled us to capture the *las* BGC minimally on 5 fosmids (Fig. 3) and to subsequently reassemble the BGC into a plasmid for heterologous expression. Network analysis with BiG-SCAPE (32) revealed no shared families with MIBiG reference BGCs indicating the novelty of the *las* BGC.

**Fig. 3.**
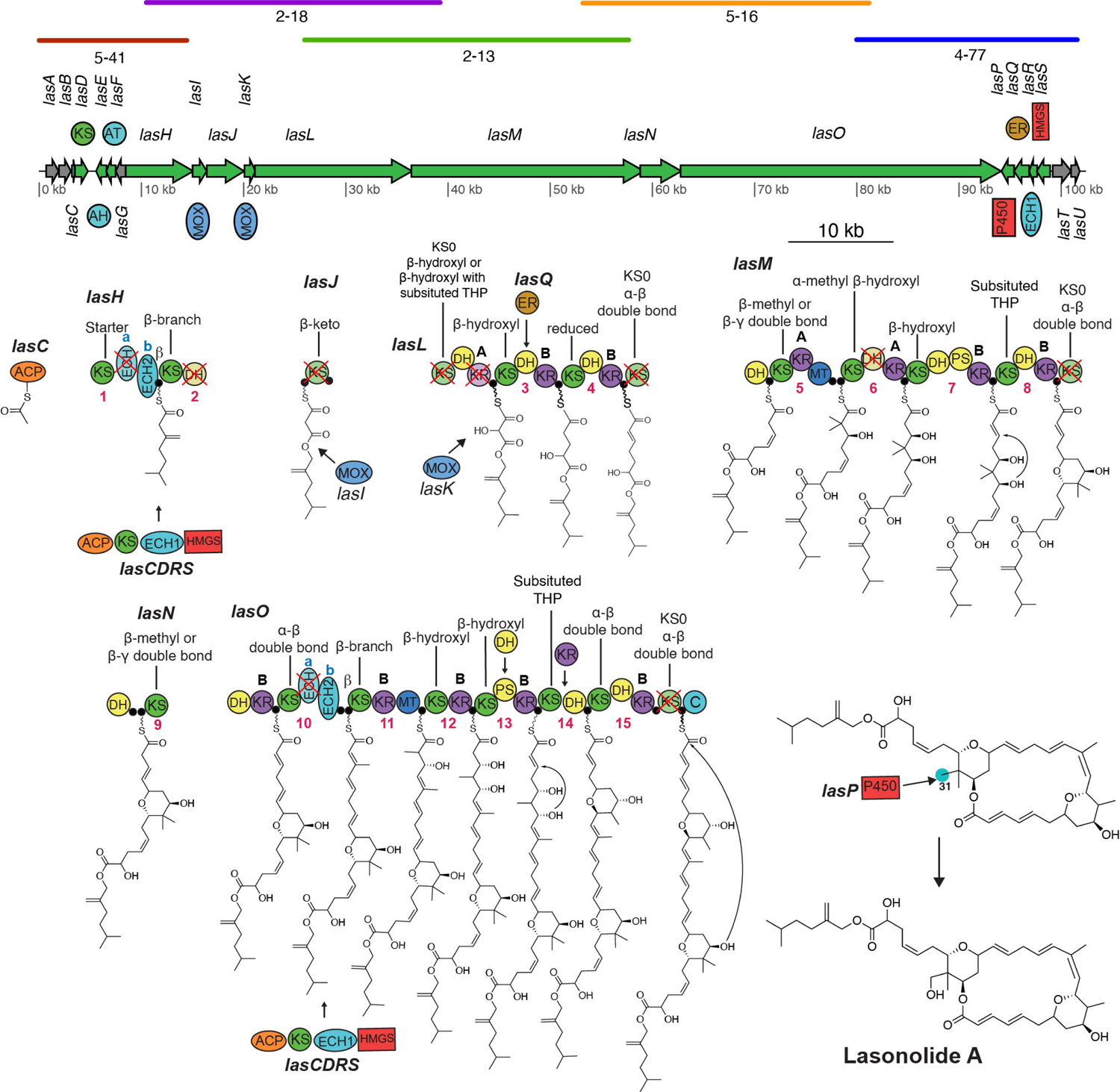
Proposed LSA biosynthetic scheme. Colored lines above the *las* BGC represent an alignment of individual fosmids to the pathway, the fosmids were subsequently assembled together in a plasmid. A cross indicates a domain predicted to be catalytically inactive. Open reading frames colored in gray represent genes with unknown or no role in LSA synthesis. Numbers below domains indicate the module number and ‘A’ and ‘B’ denote the predicted stereoconfiguration of the KR product, as previously described (50, 51). Predicted substrate specificity of KS domains, obtained through phylogeny (**Fig. S3**) (33), are shown above each respective KS domain. Carbon 31 is highlighted in blue to represent the site where P450 LasP is predicted to act. Abbreviations: ACP, acyl carrier protein, also denoted by a filled black circle; AH, acylhydrolase; AT, acyltransferase; C, condensation; DH, dehydratase; ECH, enoyl-CoA reductase; ER, enoylreductase; HMGS, 3-hydroxy-3-methylglutaryl-CoA synthase; KR, ketoreductase; KS, ketosynthase; MOX, monooxygenase; PS, pyransynthase; P450, cytochrome P450; THP, Tetrahydropyran.

### Model for lasonolide biosynthesis by *las* BGC

The proposed biosynthetic scheme for the synthesis of LSA based on the *las* BGC is shown in **Fig. 3**. The complete *las* BGC consists of six *trans*-AT PKS proteins (*lasHJLMNO*), ten accessory genes (*lasCDEFIKPQRS*) and five genes with no or unknown role in LSA synthesis (*lasABGTU*). Phylogenetic analysis of 944 different KS domains (**Fig. S2**) was used to predict KS substrate specificity (33), which was found to be similar to the proposed biosynthetic model. The pathway is predicted to be collinear with the first KS domain of *lasH* clustering into the same clade as other starter KS domains in the KS phylogenetic tree. Moreover, the last *trans-*AT PKS protein (*lasO*) contains a condensation domain, similar to those found in nonribosomal peptide synthetase pathways, as its terminal domain, which is proposed to be responsible for cyclizing and cleaving the final PKS product (24).

An acylhydrolase (AH) domain is often used in *trans*-AT PKS systems for proofreading by cleaving the acyl units from stalled sites (34, 35). AHs are closely related to acyltransferase (AT) domains, which are involved in the addition of malonyl-*S*-coenzyme A extender units on the phosphopantetheine arms on ACP domains (24, 36). LasE (AH) and LasF (AT) were correctly identified as AH and AT domains respectively, based on the presence of active site residues (**Fig. S3A**) and phylogeny (35) (**Fig. S3B**). The accessory proteins LasCDRS include enzymes known-branch formation at module 1 and 10 (21). The ACPs at module 1 and 10 contain a conserved tryptophan which is involved in interacting with β 38). LasR was identified to be responsible for dehydration (ECH1) while LasH ECHb and LasO ECHb to be responsible for decarboxylation (ECH2) during β their truncated size and lack of homology to the conserved sites needed for oxyanion hole formation LasH ECHa and LasO ECHa are proposed to be inactive (40, 41) (**Fig. S3D and S3E**). An endo-β-methyl (β-unsaturated β-methyl) is predicted to form on module 10. The presence of a truncated ECH domain just upstream of ECH2 domain has been commonly observed with the formation of exo-β-methylene (β,γ-unsaturated β-methylene), but to our knowledge this is the first time such an architecture has been reported to form a endo-β-methyl (38). Based on the collinearity of the pathway suggested above and the split module architecture (KS-DH MOX ACP-KS) associated with different Baeyer-Villiger (BV) monooxygenases as seen in oocydin and sesbanimide biosynthesis (42–44) we propose LasI to be involved in BV oxidation and LasK in the addition of a hydroxyl group. Based on the recent reports that the most common transformation by cytochrome P450 enzymes in PKS biosynthesis is C-H hydroxylation (45) we suggest LasP to be oxidizing C-31. Another accessory protein, the enoylreductase (ER) domain LasQ (46) is proposed to be acting in *trans* as observed in other pathways including lagriamide (47), patellazoles (48) and bacillaene (24, 49).

Due to the disruption of the catalytically active residues (CHH, **Fig. S3F**), we predict certain KS domains to be inactive (LasL KS1, LasL KS4, LasM KS5 and LasO KS7). We propose that the ACP domain of LasL directly takes the molecule from the first ACP of LasJ and thus we predict the KS domain in LasJ to be catalytically inactive despite the presence of catalytic residues, as observed in lagriamide, lankacidin, and etnangien pathways (24, 36). Likewise, the alignment of ketoreductase (KR) domains (**Fig. S3G**) allowed us to identify the ones lacking the KSY catalytic triad and thus spot the inactive KR domain in module 2 (LasL KR1). Additionally, it was found that the predicted stereoconfiguration of KR products (50, 51) in the *las* BGC, matched the configuration of the equivalent moieties within the LSA structure produced by total synthesis (8). The absence of a KR domain required in module 14 is proposed to be compensated by a *trans*-acting KR likely from the following module as proposed in the patellazole (48) pathway.

We were able to identify two pyran synthase (PS) domains (in module 7 and module 13) based on their phylogeny (**Fig. S4A**) and alignment (52, 53) (**Fig. S4B**). These PS domains are at the correct position in the *las* BGC to insert the pyran rings required to synthesize LSA. Even though module 13 lacks a DH domain required for pyran ring formation, we predict this role to be played by a *trans*-acting DH domain as commonly seen in *trans*-AT PKS pathways (24). Similarly, we were able to identify double bond-shifting DH domains in module 4 (LasM DH1) and 8 (LasN DH1) by the absence of both proline at the HxxxGxxxxP motif and of Glutamine/Histamine at the DxxQ/H (**Fig. S4C**) (54) motif. Moreover, alignment of the DH domains allowed us to identify the presence of inactive DH domains in module 2 (LasH DH1) and 6 (LasM DH2) by the absence of catalytic histidine at the HxxxGxxxxP motif and catalytic aspartic acid at the DxxxQ/H motif (**Fig. S4D**). LasL DH3 has a serine in place of proline in its HxxxGxxxxP motif. Alignment with different DH domains with serine in the HxxxGxxxxP motif revealed a mixture of domains annotated as active and inactive (**Fig. S4E**). The majority of times, when the DH domain had the conserved histidine in the HxxxGxxxxP motif it was annotated as active. Based on this we propose LasL DH3 to be active. Specific primers were designed based on the *las* BGC sequence and used to identify additional fosmids so that the whole pathway could be assembled from five overlapping clones for future heterologous expression.

For the biosynthesis of other LSA analogs, we propose that all of them except for lasonolide D are modified post PKS (**Fig. 4**). The cytochrome P450 LasP is predicted to oxidize LSA at C-37 and C-36 leading to the synthesis of lasonolide B and C respectively. However, in the complete biosynthesis of lasonolide B it is unclear how the methyl group is transferred from C-38 in LSA to C-36 in lasonolide B. Recently it was shown that serine hydrolase activity of lipid droplet-associated hydrolase is responsible for cleaving the ester bond in LSA and yielding the active form of the molecule, i.e. lasonolide F (55). Due to its hydrophobicity LSA is able to easily diffuse into the plasma membrane and into lipid droplets, where it is converted into lasonolide F, which is more hydrophilic and therefore able to diffuse out of the lipid droplet and into the cytoplasm to exhibit its cytotoxic effect (55). Lasonolide C seems to undergo an esterification reaction with a long-chain fatty acid (CH_3_(CH_2_)_10_COOH) to produce lasonolide G. We suggest that lasonolide E is also biosynthesized by a *trans*-esterification reaction, by reacting with an ethanol molecule. As with the production of LSA, we suggest that for the biosynthesis of lasonolide D the molecule passes through the entire *las* BGC, however, the starter molecule in this case is an acetate instead of a malonate that gets loaded on the ACP of LasJ, with LasH and LasI being inactive.

**Fig. 4.**
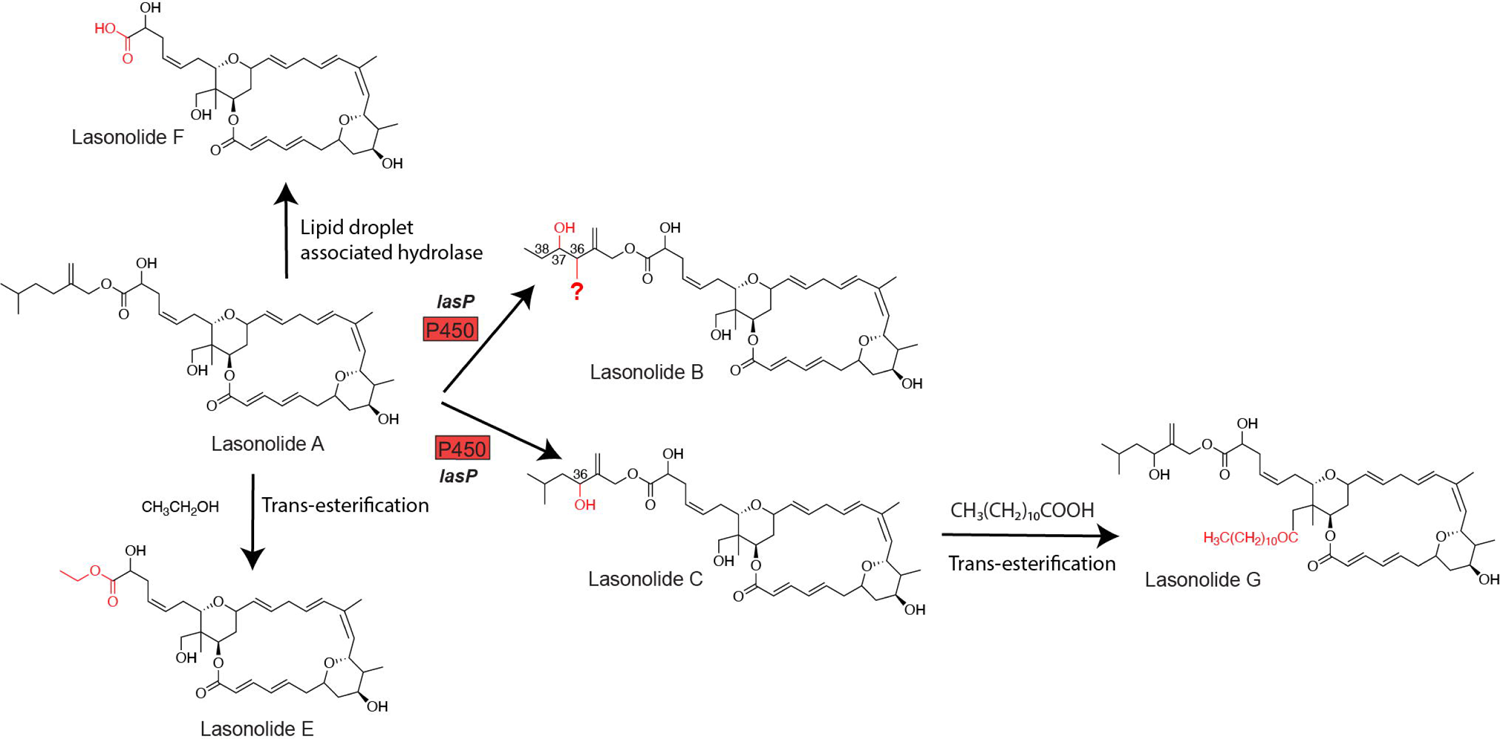
Proposed biosynthesis of different analogs from lasonolide A. We could not identify an enzyme which would explain the migration of the methyl from C-38 in LSA to C-36 in lasonolide B.

### Multiple repeats of the *las* BGC

The k-mer coverage of the *las* BGC (400.165× for *las* BGC_v1 and 159.02× for *las* BGC_v2) is roughly three times that of “*Ca.* T. lasonolidus” (135.16x in Forcepia_v1 and 48.24x in Forcepia_v2). The 3× coverage suggests three repeats of the putative *las* BGC. Visual inspection of the assembly graph as well as mapping of the paired-end reads onto “*Ca.* T. lasonolidus” allowed us to identify three connections on the 3′ end of *las* BGC but only two connections on the 5′ end of the pathway (contig 7 and 8) (**Fig. 5** and **Table S3**). Another contig (contig 5) was observed to be connected to *las* BGC about 3 kbp (3.6 kbp for *las* BGC_v1 and 3.7 kbp for *las* BGC_v2) from the 5LJ end of *las* BGC. This suggests that the majority of *las* BGC (about 98kbp) is repeated thrice with a 3 kbp segment of the pathway (contig 6) is repeated twice (**Fig. 5**). The two repeats of contig 6 were further verified by more than twice paired end reads mapping to it as compared to contig 5 (56) as well as its 2× coverage when compared to “*Ca.* T. lasonolidus”. All the connections between the *las* BGC and the bacterial genome were verified using PCR and Sanger sequencing of the amplicons. We believe that the three repeats of the *las* BGC might be involved in contributing to the increased expression of LSA through increased gene dosage (57).

**Fig. 5.**
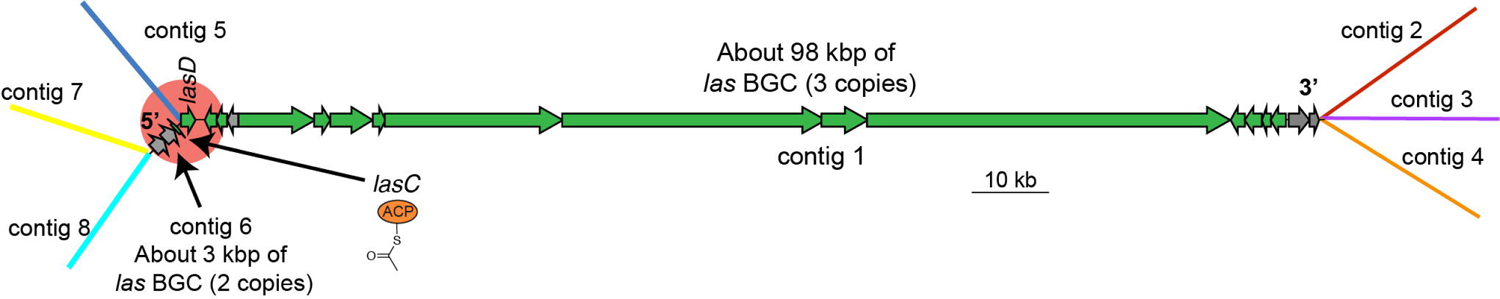
Model for three repeats of the *las* BGC. The 5′ end of *las* BGC is highlighted to demonstrate the location where one of the *las* BGC repeats lacks *lasC*. Contig(s) making up the 98 kbp segment of *las* BGC (one in *las* BGC_v1 and six in *las* BGC_v2) have been collectively referred to as contig 1. Contigs except for the *las* BGC are not shown to scale.

On comparing the three repeats it was observed that the *las* BGC repeat connected to contig 5 lacks *lasC* (ACP domain, highlighted area in **Fig. 5**), which is predicted to play an important role in β-branch formation. Furthermore, the same repeat which lacks *lasC* also shows the presence of an incomplete *lasD* (decarboxylating KS domain used in β-branching). Although this KS domain has the catalytic active site residues SHH, characteristic of decarboxylating KSs (38), it lacks about 47 amino acids that are present in the KS domain of the other two repeats connected to contig 6. On further investigation with GATK HaplotypeCaller (58, 59) we were able to detect three insertions and two single nucleotide polymorphisms (SNPs) between the three repeats of *las* BGC_v1 (**Fig. 6** and **Table 1**). This was further supported by the allelic depth (AD) - informative reads supporting each allele - and phred-scaled likelihoods (PL) of the possible genotypes. The genotype quality (GQ) which represents the confidence in the PL values was 99 for all five variants, which is the maximum value GATK reports for GQ. Furthermore, alignment of *las* BGC_v1 with *las* BGC_v2 revealed that *las* BGC_v2 contains all the variants that were called by GATK, thus further supporting their presence. All the three insertions are multiples of three base pairs (60bp, 24bp and 54bp), and are thus not causing any frame-shift mutations. Moreover, all the three insertions lie between *trans-*AT PKS domains, suggesting they do not contribute to functional differences. Change in one base from G to A at 93,995 bp does not result in a change in the amino acid sequence as both codons (TAT and TAC) encode for tyrosine. Finally, the change in base from A to G at 95,154 bp lies just outside *lasS*, i.e. in the non-coding region. The above-mentioned differences in the three repeats of *las* BGC indicate that the repeats have been present long enough to allow divergence. However, the differences between the three repeats are not predicted to affect the function of the *las* BGC.

**Fig. 6.**
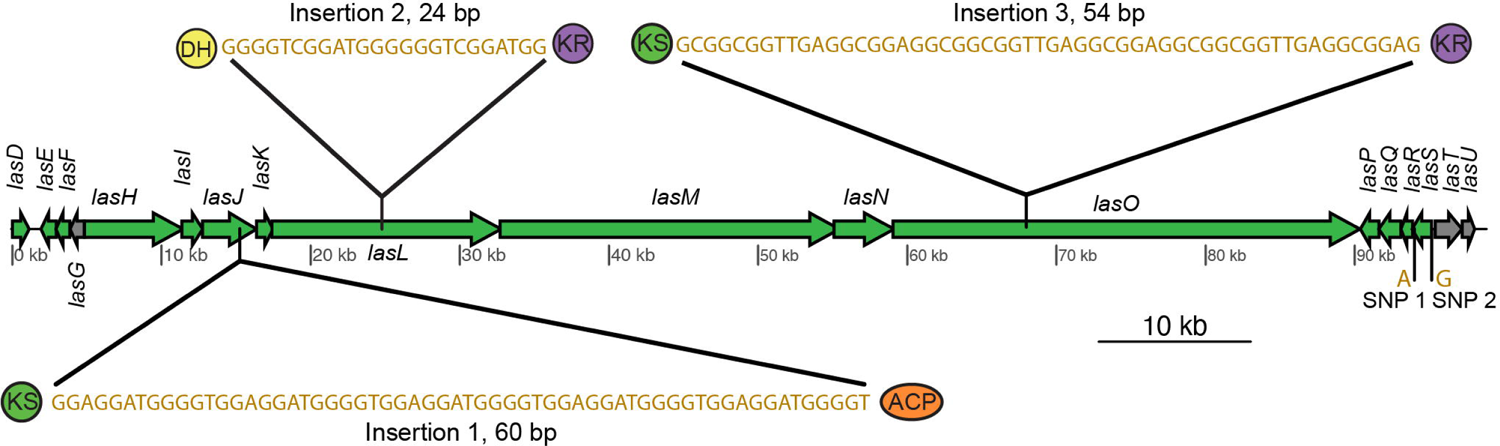
Variants identified between the three repeats of *las* BGC_v1.

**Table 1.**
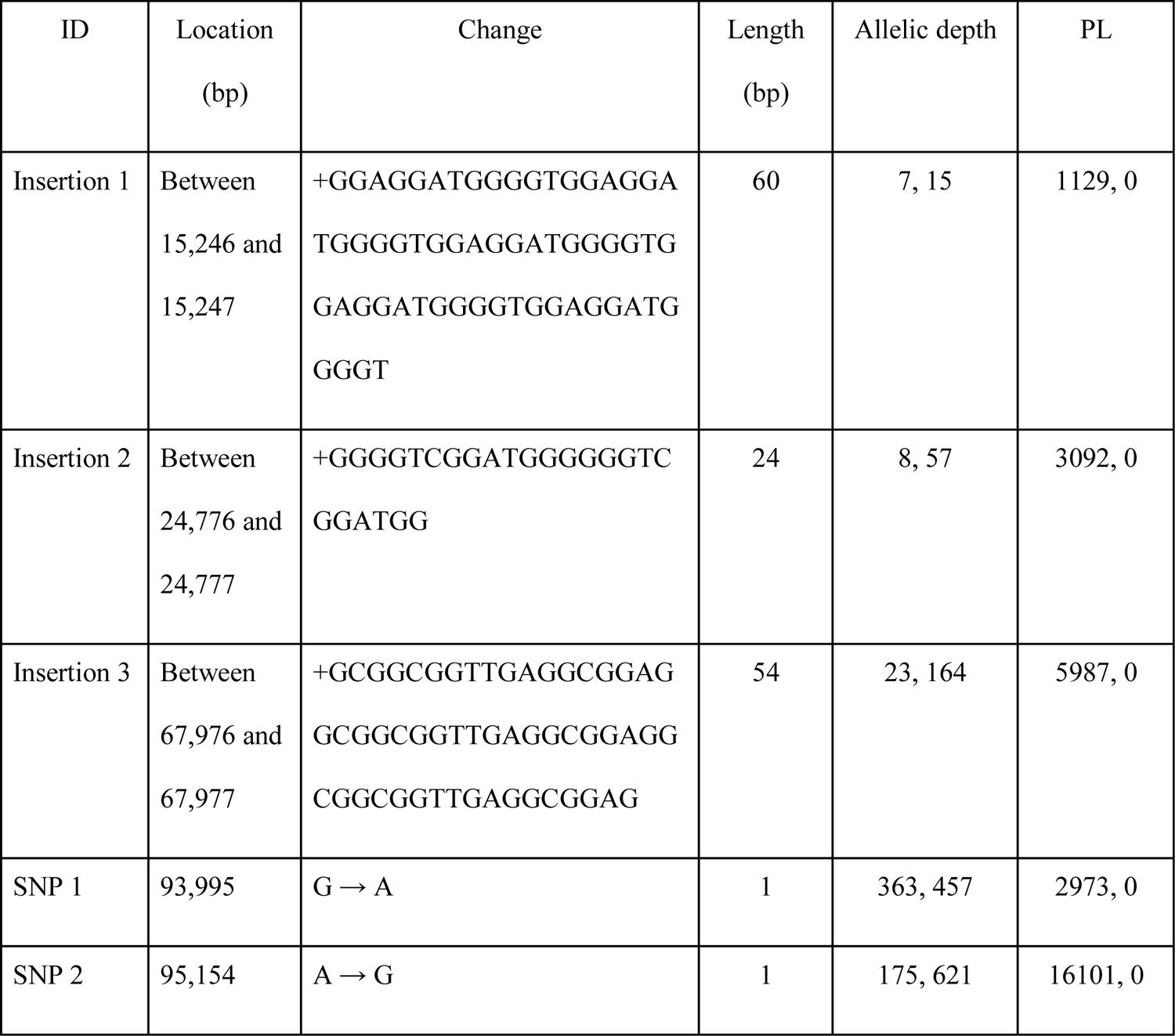
Description of the variants identified between the three repeats of *las* BGC_v1. Both AD and PL values are represented in the manner “reference, variant”. A lower PL value represents a higher likelihood of the sample being that genotype.

### Evidence for horizontal gene transfer

During the binning process by Autometa (60), Barnes-Hut Stochastic Neighbor Embedding (BH-tSNE) was used to reduce 5-mer frequencies to two dimensions. Generally, contigs belonging to the same genome would have similar 5-mer frequency and would be expected to cluster close to each other (61, 62). Visualization of the dimension-reduced data (**Fig. 7A–B and S5A–B)**, revealed that the *las* BGC contigs significantly differ in their 5-mer frequency from “*Ca.* T. lasonolidus”, suggesting that the *las* BGC could have been recently horizontally acquired. Furthermore, the GC% of the *las* BGC is significantly different (p < 0.05, ANOVA followed by Tukey HSD) from annotated, hypothetical and pseudogenes (**Fig. 7C and S5C**) providing further evidence for horizontal transfer of the *las* BGC.

**Fig. 7.**
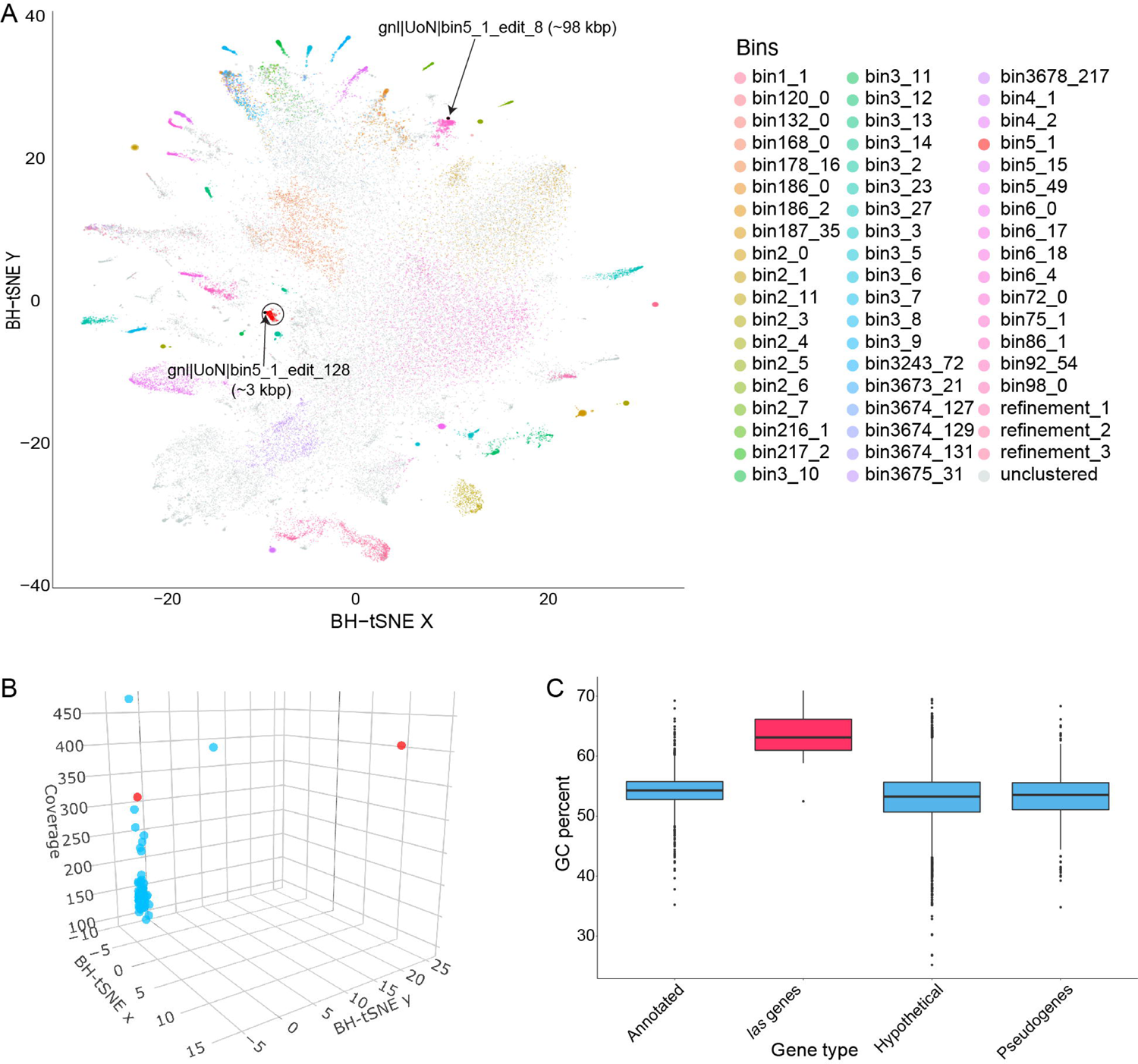
**(A)** 2D visualization of Automata binning of Forcepia_v1. The “*Ca.* T. lasonolidus” genome is circled in black and the *las* BGC contigs are marked with an arrow. Axes represent dimension-reduced Barnes-Hut Stochastic Neighbor Embedding (BH-tSNE) values (BH-tSNE x and BH-tSNE y). **(B)** 3D visualization of contigs present in “*Ca.* T. lasonolidus”. The *las* BGC is colored red. Axes represent BH-tSNE values (BH-tSNE x and BH-tSNE y) along with k-mer coverage. **(C)** GC percentage of different sets of genes in Forcepia_v1 “*Ca.* T. lasonolidus”. The *las* BGC genes are colored in red.

Codon adaptation index (CAI) compares the synonymous codon usage of a gene and that of a reference set along with measuring the synonymous codon usage bias (63). The CAI for the *las* BGC was significantly different (p < 0.05, ANOVA followed by Tukey HSD) from the annotated, hypothetical and pseudogenes, but matched that of highly expressed genes (i.e. ribosomal proteins) (**Fig. S5D-E**). Thus, despite its horizontal acquisition, the BGC’s codon usage has been adapted to be efficiently translated even though the 5-mer composition is still different when compared to the rest of the “*Ca.* T. lasonolidus” genome.

### The genome of the putative lasonolide producing symbiont

“*Ca.* T. lasonolidus”, with multiple *las* BGC repeats, represents an important addition to the growing collection of symbiotic Verrucomicrobia (“*Candidatus* Didemnitutus mandela’’ and “*Candidatus* Synoicihabitans palmerolidicus”) being identified with repeated *trans*-AT PKS BGCs (57, 64, 65). Recently, two simultaneous studies have also identified a *trans*-AT PKS BGC for pateamine in a bacterium (“*Candidatus* Patea custodiens”) belonging to phylum Kiritimatiellaeota (66, 67), a recently proposed phylum which was previously classified within Verrucomicrobia (68). This highlights the importance of this understudied phylum as an important producer of natural products. “*Ca.* T. lasonolidus” is little over 5 Mbp long and has GC percentage of about 53%. It is estimated to be 99% complete, is 1.35% contaminated (69), has tRNAs for all amino acids and complete 5S, 16S and 23S rRNA genes. Based on MIMAG standards (25) the bin is classified as a high-quality MAG. Detailed statistics of the putative LSA producer are provided in **Table 2**.

**Table 2.**
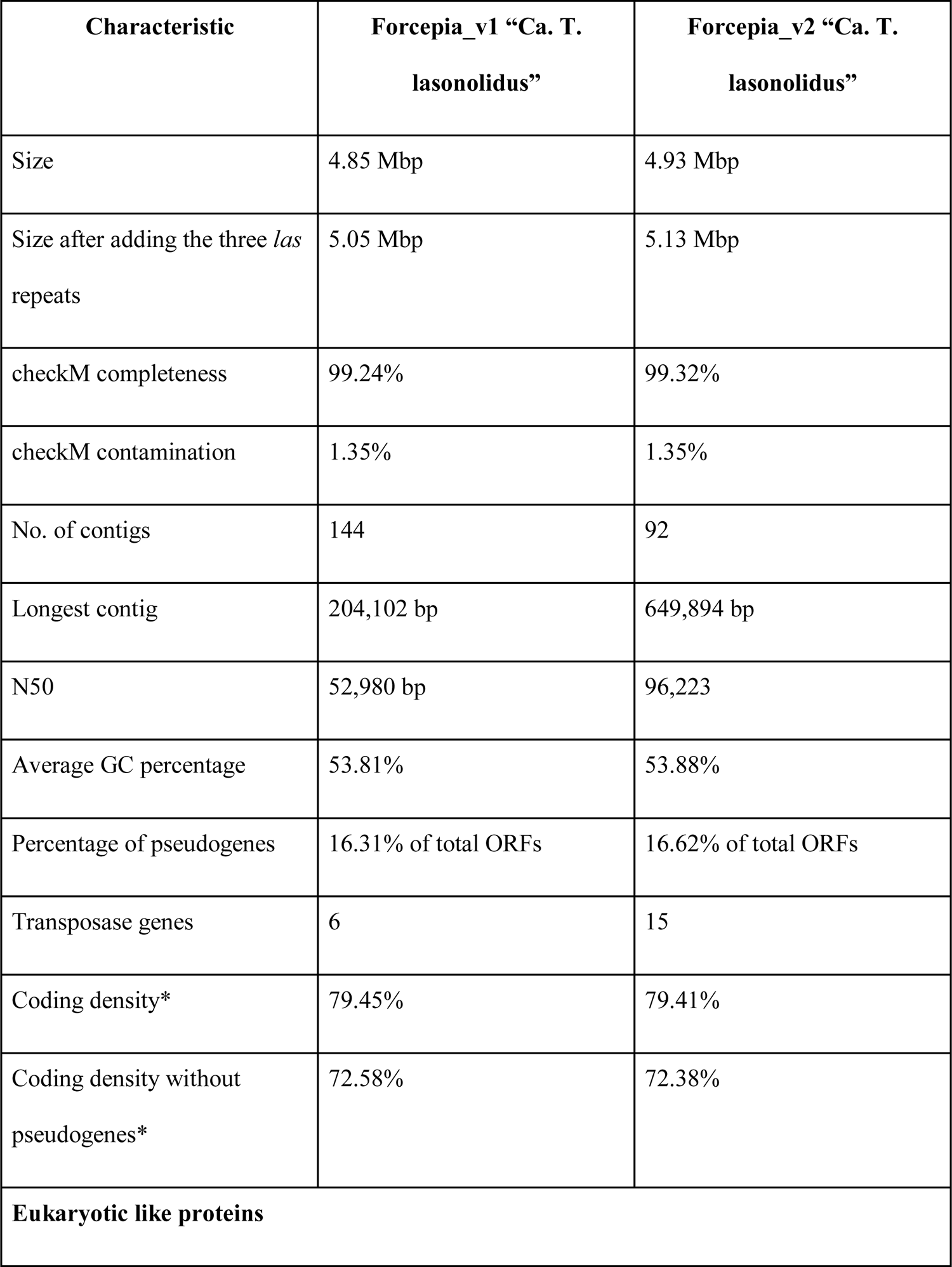
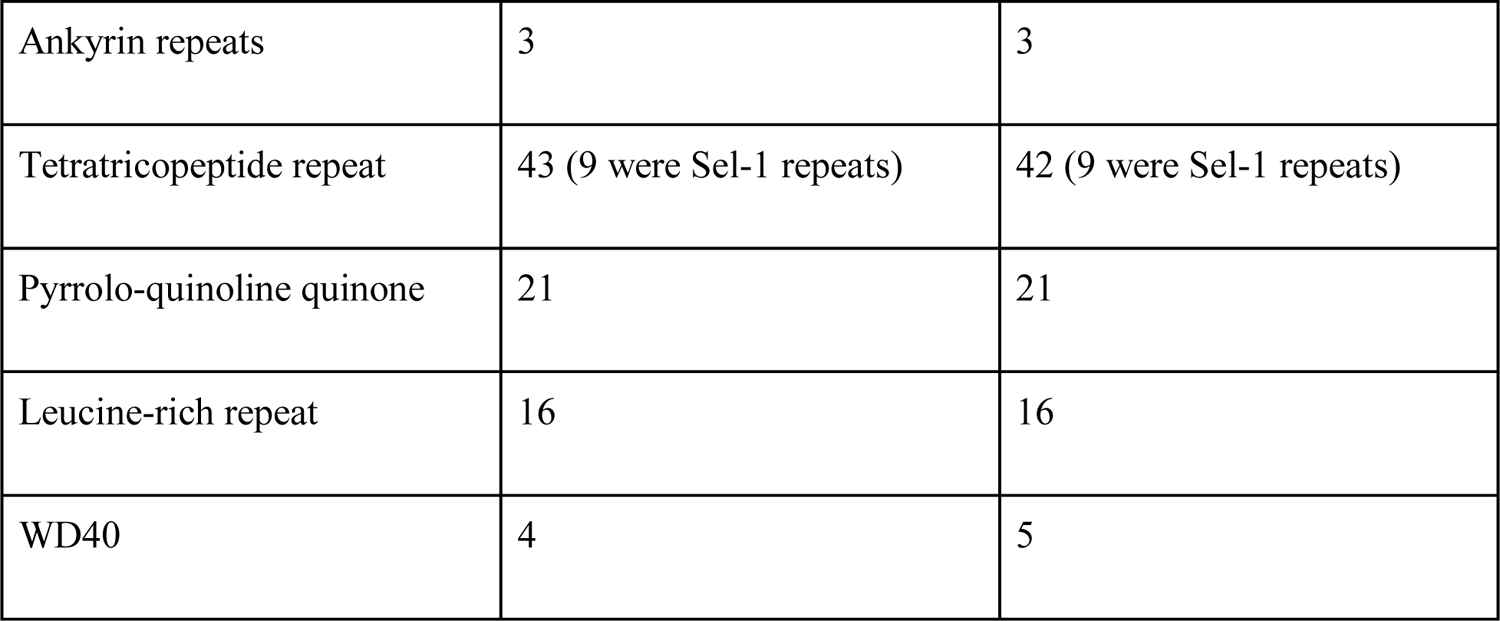
Genome statistics for “*Ca.* T. lasonolidus”. *Coding density is weighted by length taking into account the 97.11% coding density of *las* BGC repeats.

Eukaryotic-like proteins (ELPs) are known to be present in genomes of sponge symbionts and have been found to play an important role in regulating their interaction with the host sponge (70–73). It is hypothesized that interaction with ELPs allow the symbiotic bacteria to evade phagocytosis by the sponge, thus allowing discrimination between food and symbiont bacteria (72, 74). A number of ELPs were identified in “*Ca.* T. lasonolidus” (**Table 3, and Table S4A**), thus suggesting a symbiotic relationship of the bacterium with the *Forcepia* sp.

Bacterial microcompartments (BMCs) are organelles that enclose enzymes within a selectively permeable proteinaceous shell (75), and they are rare among bacteria. Members of the phyla Planctomycetes and Verrucomicrobia have a unique BMC gene cluster called the Planctomycetes-Verrucomicrobia bacterial microcompartment (PV BMC) which is responsible for production of microcompartment shell proteins BMC-P and BMC-H, as well as degradation of L-rhamnose, L-fucose and fucoidans (71, 76, 77). Genes encoding the PV BMC cluster were identified in “*Ca.* T. lasonolidus” (**Table S4B**), and the respective gene clusters in Forcepia_v1 “*Ca.* T. lasonolidus” and Forcepia_v2 “*Ca.* T. lasonolidus” were compared using clinker (28) and were found to be 100% identical to each other. One interesting finding was that the identified PV BMC clusters had a DNA-methyltransferase and PVUII endonuclease gene between the first and the second BMC-H genes. This is different from the usual arrangement of the PV BMC gene cluster where both the BMC-H genes lie next to each other and the cluster lacks DNA-methyltransferase and PVUII endonuclease genes (**Fig. 8**). The presence of PV BMC genes in the “*Ca.* T. lasonolidus” genome suggests that it possesses bacterial microcompartments and that they might be involved in L-fucose and L-rhamnose degradation. Despite repeated attempts, we only found rhamnulokinase and fumarylacetoacetate hydrolase family proteins in the “*Ca*. T. lasonolidus” genomes and failed to identify other complementary enzymes involved in the degradation of L-fucose and L-rhamnose. However, other enzymes involved in carbohydrate metabolism including glycoside hydrolases, carbohydrate binding module, polysaccharide lyase, carbohydrate esterases and glycoside transferase were detected (**Table S4C**) indicating that “*Ca.* T. lasonolidus” is capable of polysaccharide degradation, something that is observed in a number of marine Verrucomicrobia (78–80).

**Fig. 8.**
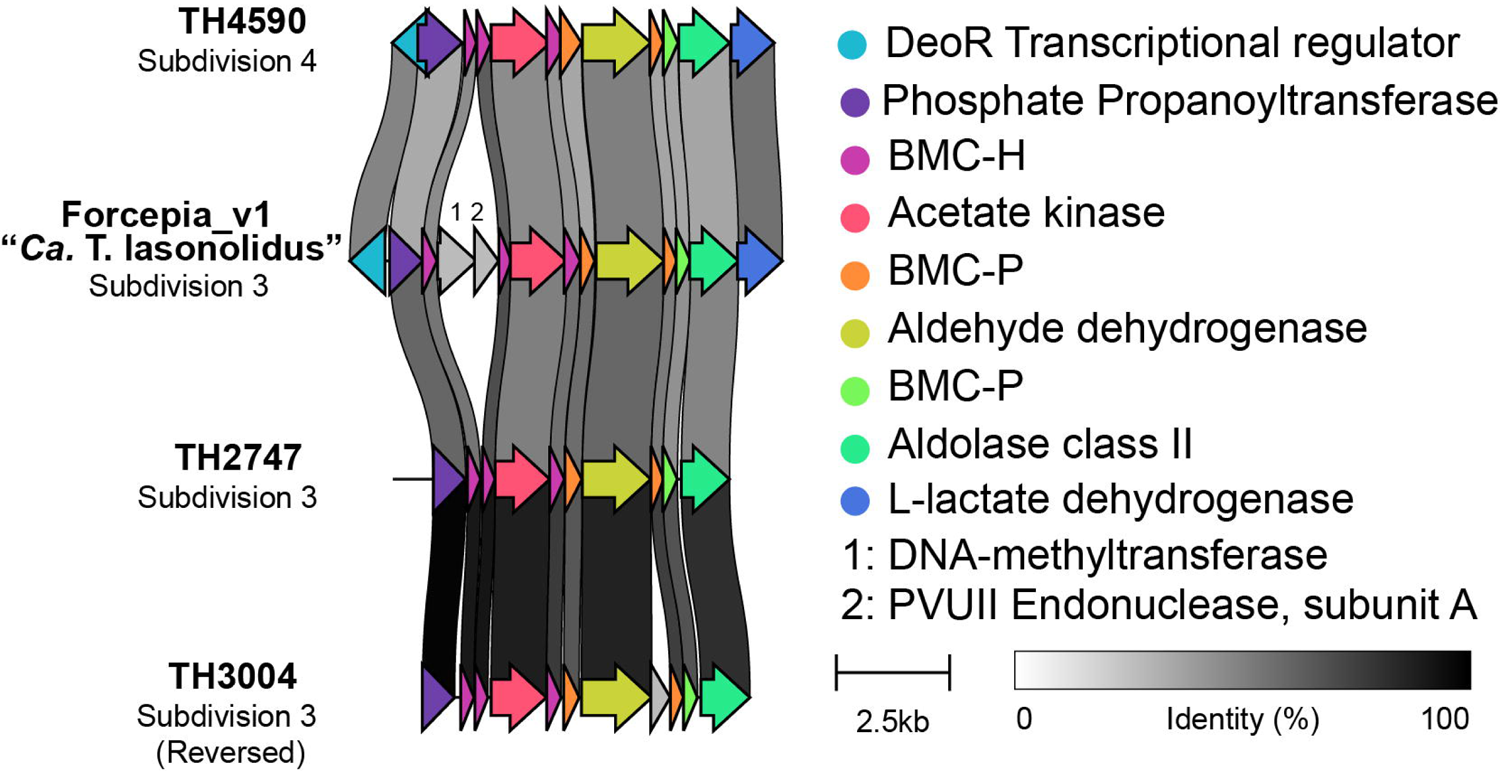
Comparison of PV BMC gene cluster in Forcepia_v1 “*Ca.* T. lasonolidus” with the PV BMC cluster from other Verrucomicrobia. “*Ca.* T. lasonolidus” has DNA-methyltransferase and PVUII endonuclease genes (in gray, labeled 1 and 2) between the first and the second BMC-H genes. This kind of arrangement was not observed in other PV BMC clusters.

A characteristic of obligate host-symbiont relationships is the loss of symbiont genes which are required for independent survival. The early stages of genome reduction are characterized by reduced coding density, and a high number of pseudogenes (81–83). We compared “*Ca* T. lasonolidus” with its closest free-living relative - *Pedosphaera parvula* Ellin514 (GCA_000172555.1). The draft genome of *P. paruva* Ellin514 is 7.41Mbp long, about 2.2 Mbp longer than “*Ca.* T. lasonolidus”. Furthermore, in *P. paruva* Ellin514 only 0.5% of total ORFs were found to be pseudogenes (57, 84, 85) as opposed to about 16% in “*Ca.* T. lasonolidus” (**Fig. 9A-B**). Another indication of ongoing genome reduction comes from the fact that a much smaller percentage of genes with annotated function were identified in “*Ca.* T. lasonolidus” as compared to *P. paruva* Ellin514 (**Fig. 9C**), perhaps indicating sequence degradation and divergence from functionally-annotated genes. Moreover, when compared with *P. paruva* Ellin514, “*Ca.* T. lasonolidus” lacks genes involved in DNA repair, DNA replication, chemotaxis and nucleotide metabolism (**Fig. 9D**), a trend which is commonly observed in symbionts undergoing genome reduction (81). However, “*Ca.* T. lasonolidus” contains most of the primary metabolic pathways (**Fig. 9E**) when compared to *P. paruva* Ellin514, and has a fairly large genome to be classified as reduced. Based on the above evidence we suggest that “*Ca.* T. lasonolidus” is in early stages of genome reduction. This hypothesis is also supported by its low coding density of ∼72% (without pseudogenes), relative to the average coding density of 85-90% for free-living bacteria (81) which suggests a recent transitional event, such as host restriction (81).

**Fig. 9.**
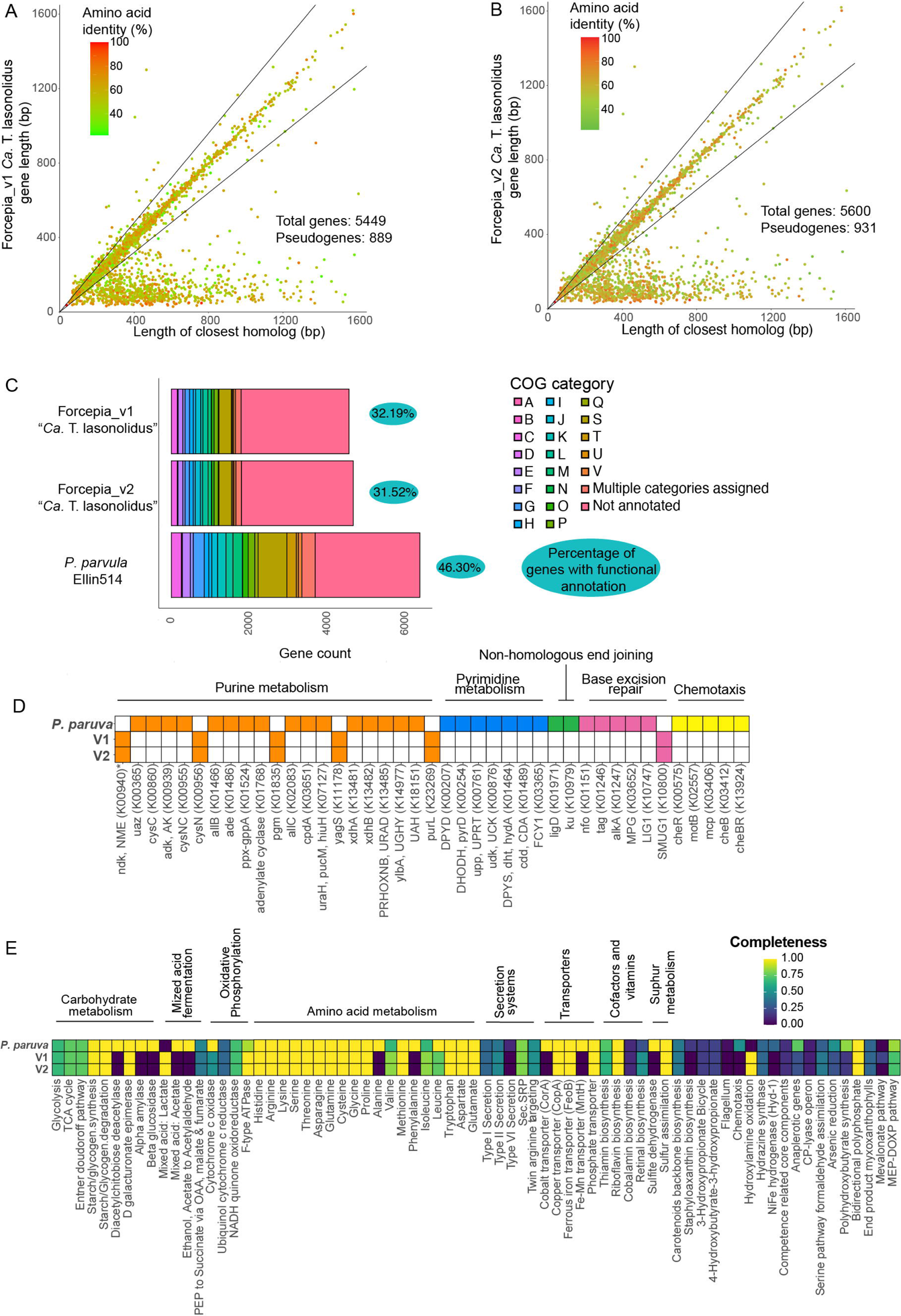
**(A and B)** Comparison of the gene length in **(A)** Forcepia_v1 “*Ca.* T. lasonolidus” and **(B)** Forcepia_v2 “*Ca.* T. lasonolidus”, respectively with their closest homologs in the nr database. Genes with length less than 80% of the closest homolog (below the lower black line) are classified as putative pseudogenes (57, 84, 85). The graphs have been truncated for clarity, as some genes are many kbp long. **(C)** Comparison of functional COG categories in Forcepia_v1 “*Ca.* T. lasonolidus”, Forcepia_v2 “*Ca.* T. lasonolidus” and *P. paruva* Ellin514 for non-pseudogenes. A gene is considered to have a functional annotation when it belongs to a COG category, except for category S which represents unknown function. **(D)** Comparison of genes in different metabolic pathways for Forcepia_v1 “*Ca.* T. lasonolidus”, Forcepia_v2 “*Ca.* T. lasonolidus” and *P. paruva* Ellin514, including only non-pseudogenes. Colored squares represent presence of a gene while white squares represent absence of gene. *K00940 is involved in both purine and pyrimidine metabolism. Genes absent in all three genomes have been removed. **(E)** Comparison of completeness of different metabolic pathways in Forcepia_v1 “*Ca.* T. lasonolidus”, Forcepia_v2 “*Ca.* T. lasonolidus” and *P. paruva* Ellin514 (including only non-pseudogenes) as determined by KEGG decoder (86). Pathways have been grouped into categories wherever possible. Pathways absent in all three genomes have been removed. V1 and V2 refer to Forcepia_v1 “*Ca.* T. lasonolidus” and Forcepia_v2 “*Ca.* T. lasonolidus” respectively.

Due to its potency and unique mechanism of action, LSA is considered a potential anti-cancer drug lead; however, its limited supply has hampered its transition to clinical trials. The evidence provided here suggests that LSA is synthesized by a yet uncultured Verrucomicrobial symbiont, which harbors three copies of the putative *las* BGC. The detailed analysis of the biosynthetic scheme, genome characteristics of the putative producer as well as the capture of the *las* BGC on a plasmid will aid future cultivation and heterologous expression efforts.

## Acknowledgments

The authors acknowledge Amy Wright and Shirley Pomponi for providing sponge specimens, and Amy Wright, Shirley Pomponi and Peter McCarthy for valuable discussions during the project. The authors also thank Samantha C. Waterworth for fruitful discussions, John Barkei for discussions on heterologous expression strategies, and Chase Clark for providing DH sequences for sequence comparisons.

## Data availability

The data associated with this study is deposited under BioProject RJNA833117. The WGS reads have been deposited in the sequence read archive (SRA) with accessions SRR18966768 (Forcepia_v1) and SRR18966767 (Forcepia_v2).

## Methods

For full details see **Text S1.**

### Sponge collection

*Forcepia sp.* (class, *Demospongiae*; order, *Poecilosclerida*; family, *Coelosphaeridae*) was collected in August of 2005 using the HBOI Johnson Sea Link submersible. Samples were collected at a depth of 70m from the Gulf of Mexico (26.256573N, 83.702772W) on the Pulley Ridge (https://shinyapps.fau.edu/app/bmr). The sponge samples were immediately frozen at −80°C. The sample ID was 12-VIII-05-1-006 200508121006 2005-08-12 JSL I-4837 (HBOI) *Forcepia* sp. 131921.

### DNA purification and sequencing

The sponge hologenome was extracted using a modified cetyl trimethylammonium bromide (CTAB) DNA extraction method (87) and then size-fractionated by low melting point gel electrophoresis. DNA fragments greater than 40 kb were recovered from the gel and used for fosmid library preparation (**Text S1**) as well as metagenomic sequencing. Two rounds of sequencing were performed for different DNA extracts from the *Forcepia* sp. sponge. For the first round (referred to as Forcepia_v1) Illumina TruSeq DNA libraries were prepared and sequenced by RTL Genomics using an Illumina MiSeq sequencer giving us 108 million paired-end reads with length of 151bp. For the second round of sequencing (referred to as Forcepia_v2) Illumina Nextera libraries were prepared and sequenced using a NovaSeq 6000 sequencer giving us 303 million paired-end reads with length of 150 bp. Fosmids were sequenced by RTL Genomics and Genewiz.

### Identification and annotation of *las* BGC

Identification of the *las* BGC was done using tBLASTN (26), where KS domains from different *trans*-AT PKS pathways were used as a query against the metagenomic assembly (assembled using MetaSpades (88), see **Text S1**). Genes for each bin were called and annotated using Prokka v1 (89, 90). MetaSpades contig headers were replaced by their respective Prokka headers to maintain consistency with the annotation file submitted to NCBI. Genes on contigs making up the *las* BGC were not called correctly by Prokka (89, 90) and were thus annotated manually in Artemis (91) with the help of AntiSMASH (27), CDD (92) and SMART (93, 94).

### Functional analysis of the “*Ca.* T. lasonolidus” genome

Genes called using Prokka v1 were used for all functional analysis (89, 90). PV-BMC clusters were identified in “*Ca.* T. lasonolidus” using Interproscan v5.52-86.0 (95) and CDD (92). Initial identification of ELPs was done using Diamond BLASTP against the diamond-formatted nr database (using -k 1 --max-hsps 1 options) (96) and Interproscan v5.52-86.0 (95). This was followed by verification of non-pseudogenes using CDD (92). Enzymes involved in carbohydrate metabolism were detected using dbCAN2 (97) where genes annotated by ≥2 tools (out of HMMER, Diamond and Hotpep) were kept. COG categories were identified using the eggNOG mapper online server (98, 99).

The genome of *P. paruva* Ellin514 was downloaded from Genbank (GCA_000172555.1) and genes were called and annotated using Prokka v1 (89, 90). Primary metabolic pathways were identified for non-pseudogenes with kofamscan using the --mapper flag (100) and annotated against the KEGG database (101–103). The matrix with presence/absence of different enzymes was constructed in RStudio (104). Completeness of metabolic pathways was identified using KEGG-Decoder (86).

## Legend for supplementary material

**Fig S1. (A)** Alignment of fosmids to the *las* BGC. Fosmids are depicted as arrows above the *las* BGC. Fosmids captured before WGS are colored orange (3-46, 5-16, 6-17, 4-77 and 1-80), whereas fosmids captured after WGS are colored blue (5-41, 2-18, and 2-13). **(B)** Relative abundance of different phyla in the sequenced Forcepia_v2 metagenome. Each block shows the relative abundance of each metagenome-assembled genome (MAG), with different colors representing the phylum they belong to. The *las* BGC-carrying bin is highlighted is marked with a star. **(C)** Assembly graph of *las* BGC_v1 visualized in BANDAGE (Wick RR, Schultz MB, Zobel J, Holt KE, Bioinformatics 31:3350–3352, 2015, https://doi.org/10.1093/bioinformatics/btv383). **(D)** Mapping of paired-end reads to contigs making up *las* BGC_v1. Contigs in green boxes represent the *las* BGC, red boxes represent the 5′ end of *las* BGC and blue boxes represent the 3′ end of *las* BGC. **(E)** Assembly of the seven contigs making up *las* BGC_v2. **(F)** Assembly graph of *las* BGC_v2 visualized in BANDAGE (Wick RR, Schultz MB, Zobel J, Holt KE, Bioinformatics 31:3350–3352, 2015, https://doi.org/10.1093/bioinformatics/btv383). **(G)** Mapping of paired-end reads to *las* BGC_v2. Contigs in green boxes represent the *las* BGC, red boxes represent the 5LJ end of *las* BGC and blue boxes represent the 3LJ end of *las* BGC. Panels D, E, G, and H were edited for clarity by removing contigs which had either very few paired-end read connections, were mapping to themselves or were very small. **(H)** BGC distribution in Forcepia_v2 sp. Metagenome. AntiSMASH (Blin K, Shaw S, Steinke K, Villebro R, Ziemert N, Lee SY, Medema MH, Weber T, Nucleic Acid Res 47:W81–W87, 2019, https://doi.org/10.1093/nar/gkz310) annotations of bacterial contigs greater than 3000bp are shown. Each bar indicates a MAG, grouped by phylum. The star represents the MAG containing the *las* BGC. BGC annotations have been simplified into polyketide synthase (PKS), Type 1 PKS, Type 3 PKS, *trans-*AT PKS, nonribosomal peptide synthetase (NRPS), ribosomally synthesized, post-translationally modified peptide (RiPP), hgIE-KS, hgIE-KS-T1PKS, terpenes, and others. **(I)** Phylogenetic tree of 51 different Verrucomicrobia genomes. Bootstrap values were calculated using RaxML with 1000 bootstrap replicates.

**Fig S2.** Clades from a phylogenetic tree of 944 KS domains from *trans-*AT PKS and the erythromycin BGC as an outgroup, containing KS domains from the *las* BGC. Color within the individual clades corresponds to the chemical structure shown on its right. ‘i’ and ‘c’ in *lasD* KS1 indicate the incomplete and complete KS domain in *lasD* respectively.

**Fig S3. (A)** Alignment of *las* AT and AH domains with AT and AH domains from different *trans-*AT PKS pathways. Active sites as well as sites distinguishing AT and AH domains (Jenner M, Afonso JP, Kohlhaas C, Karbaum P, Frank S, Piel J, Oldham NJ, Chem Commun 52:5262– 5265, 2016, https://doi.org/10.1039/C6CC01453D) have been marked. **(B)** Phylogenetic tree of AT and AH domains. The different types of domain separate into different clades (Jenner M, Afonso JP, Kohlhaas C, Karbaum P, Frank S, Piel J, Oldham NJ, Chem Commun 52:5262–5265, 2016, https://doi.org/10.1039/C6CC01453D). **(C)** Phylogenetic tree of ECH1 and ECH2 domains. Both the domains separate into different clades (Slocum ST, Lowell AN, Tripathi A, Shende VV, Smith JL, Sherman DH, Methods Enzymol 604:207–236, 2018, https://doi.org/10.1016/bs.mie.2018.01.034). **(D-E)** Alignment of *las* **(D)** ECH1 and **(E)** ECH2 domains with respective ECH domains from other PKS pathways. Sequence that is required for the formation of the oxyanion hole which stabilizes the enolate anions is marked. LasH_a and LasO_a are proposed to be inactive as they are truncated and show poor homology to the rest of the ECH domains (Gu L, Jia J, Liu H, Håkansson K, Gerwick WH, Sherman DH, J Am Chem Soc 128:9014–9015, 2006, http://doi.org/10.1021/ja0626382, Matilla MA, Stöckmann H, Leeper FJ, Salmond GPC, J Biol Chem 287:39125–39138, 2012, https://doi.org/10.1074/jbc.M112.401026). **(F)** Alignment of *las* KS domains with active site (CHH) marked. ‘i’ and ‘c’ in the LasD KS indicate the incomplete and complete KS domain in different repeats of LasD respectively. LasD is a decarboxylating KS which are known to lack the active site cysteine (Walker PD, Weir ANM, Willis CL, Crump MP, Nat Prod Rep 38:723– 756, 2021, https://doi.org/10.1039/D0NP00045K). **(G)** Alignment of *las* KR domains with two from the erythromycin BGC to allow comparison. Active site residues and conserved motifs are marked. The presence or absence of the second aspartate in the LDD motif is supposed to predict the stereochemistry of the hydroxyl group (Keatinge-Clay AT, Chem Biol 14:898–908, 2007, https://doi.org/10.1016/j.chembiol.2007.07.009, Caffrey P, ChemBioChem 4:654–657, 2003, https://doi.org/10.1002/cbic.200300581). Figures have been truncated for clarity and to show only the relevant sites. In phylogenetic trees, *las* BGC domains are highlighted in white.

**Fig S4. (A)** Phylogenetic tree of DH and PS domains, which separate into different clades (Wagner DT, Zhang Z, Meoded RA, Cepeda AJ, Piel J, Keatinge-Clay AT, ACS Chem Biol 13:975–983, 2018, http://doi.org/10.1021/acschembio.8b00049). *Las* BGC DH/PS domains are highlighted in white. **(B)** Alignment of PS domains identified in the *las* BGC with PS domains from other *trans*-AT PKS pathways. The DH domain from the erythromycin BGC is used for comparison. LasO DH2 and LasM DH4 are annotated as putative PS domains. Generally, PS domains have a Hx_4_P motif instead of a Hx_8_P and they lack the catalytic aspartate at the DxxxQ/H motif (Wagner DT, Zhang Z, Meoded RA, Cepeda AJ, Piel J, Keatinge-Clay AT, ACS Chem Biol 13:975–983, 2018, http://doi.org/10.1021/acschembio.8b00049, Pöplau P, Frank S, Morinaka BI, Piel J, Angew Chem Int Ed Engl 52:13215–13218, 2013, https://doi.org/10.1002/anie.201307406). This was found to be true only for LasM DH4 and not LasO DH2. However, identical variations from a traditional PS domain architecture are also seen in PS domains found in the mandelalide pathway (MndC DH3 and MndD DH3) (Lopera J, Miller IJ, McPhail KL, Kwan JC, mSystems 2:e00096–17, 2017, https://doi.org/10.1128/mSystems.00096-17). **(C)** Alignment of double bond-shifting DH domains identified in *las* BGC with similar domains found in other *trans*-AT PKS pathways. The DH domain from the erythromycin BGC is used for comparison. LasM DH1 and LasN DH1 are annotated as putative double bond-shifting DH domains. Generally, in DH shifting domains the conserved proline (P) in Hx_8_P motif is often replaced by either valine (V) or leucine (L). In the case of LasM DH1, a methionine (M) instead of V or L appears in the place of P, which is in line with what is observed in difficidin biosynthesis as well (Chen X-H, Vater J, Piel J, Franke P, Scholz R, Schneider K, Koumoutsi A, Hitzeroth G, Grammel N, Strittmatter AW, Gottschalk G, Süssmuth RD, Borriss R, J Bacteriol 188:4024–4036, 2006, https://doi.org/10.1128/JB.00052-06). Furthermore, DH shifting domains are sometimes characterized by the replacement of the conserved aspartic acid (D) with asparagine (N) and substitution of glutamine (Q) or histidine (H) with V or L in the DxxxQ/H motif. Even though LasN DH1 has an N in place of D in the DxxxQ/H motif, it substitutes Q/H with a serine (S). This is unusual and not found in any other double bond shifting DH. **(D)** Alignment of DH domains present in the *las* BGC with the DH domain from the erythromycin BGC. Putative PS and double bond-shifting DH domains have been excluded. LasH DH1 and LasM DH2 are annotated as inactive domains due to disrupted catalytic motifs Hx_8_P and DxxxQ/H. Even though in LasL DH1 the catalytic aspartic acid is replaced by glutamic acid (DxxxQ/H motif), we propose it is active, as a similar mutation is observed in the palmerolide BGC (Avalon NE, Murray AE, Daligault HE, Lo C-C, Davenport KW, Dichosa AEK, Chain PSG, Baker BJ, Front Chem 9, 2021, https://doi.org/10.3389/fchem.2021.802574). € Alignment of LasL DH3 with other DH domains having a serine in place of proline in Hx_8_P motif. The DH domain from the erythromycin BGC is used for comparison. Sequence headers in blue represent DH domains annotated as active while in red represent the ones annotated as inactive.

**Fig S5. (A)** 2D visualization of the initial Autometa binning of Forcepia_v2. The “*Ca.* T. lasonolidus” genome is circled in black and contigs making up the *las* BGC are marked with arrows. Axes represent dimension-reduced Barnes-Hut Stochastic Neighbor Embedding (BH-tSNE) values (BH-tSNE x and BH-tSNE y). **(B)** 3D visualization of contigs present in the “*Ca.* T. lasonolidus” genome. Contigs making up the *las* BGC are colored red. **(C)** GC percentage of different sets of genes in Forcepia_v2 “*Ca.* T. lasonolidus”. *Las* BGC genes are colored in red. **(D)** and **(E)** Codon adaptation index (CAI) of different categories of genes present in Forcepia_v1 “*Ca.* T. lasonolidus” and Forcepia_v2 “*Ca.* T. lasonolidus”, respectively. *Las* BGC genes are colored in red. P values for pairwise comparison between different categories of genes are shown in the matrix below their respective plots. Values with p < 0.05 are considered significant. Other non-significant p values are colored red. Annotated and hypothetical genes represent the genes annotated with a function and genes annotated as hypothetical respectively by Prokka (Seemann T, Bioinformatics 30:2068–2069, 2014, https://doi.org/10.1093/bioinformatics/btu153).

**Table S1.** List of oligonucleotide primers used for different purposes. **(A)** Primers used for screening the *Forcepia* sp. fosmid library before WGS. (**B**) Primers used for screening the *Forcepia* sp. fosmid library after WGS. **(C)** Primers used for confirming the presence of terminal connections with the *las* BGC.

**Table S2.** Metadata and taxonomic classification of all the MAGs

**Table S3.** Contigs making up the three repeats of the *las* BGC in “*Ca*. T. lasonolidus” in Forcepia_v1 and Forcepia_v2. Contig IDs represent the labels in **Fig. 5** of the main text.

**Table S4.** Gene annotation in Forcepia_v1 and Forcepia_v2 “*Ca.* T. lasonolidus”. **(A)** Non-pseudogenes annotated as Eukaryotic-like proteins. **(B)** Genes forming the PV BMC cluster. **(C)** Non-pseudogenes annotated by dbCAN2

**Text S1.** Supplementary methods

