## Supplementary figures and images for "Uncovering Lasonolide A biosynthesis using genome-resolved metagenomics"

### Figure S1

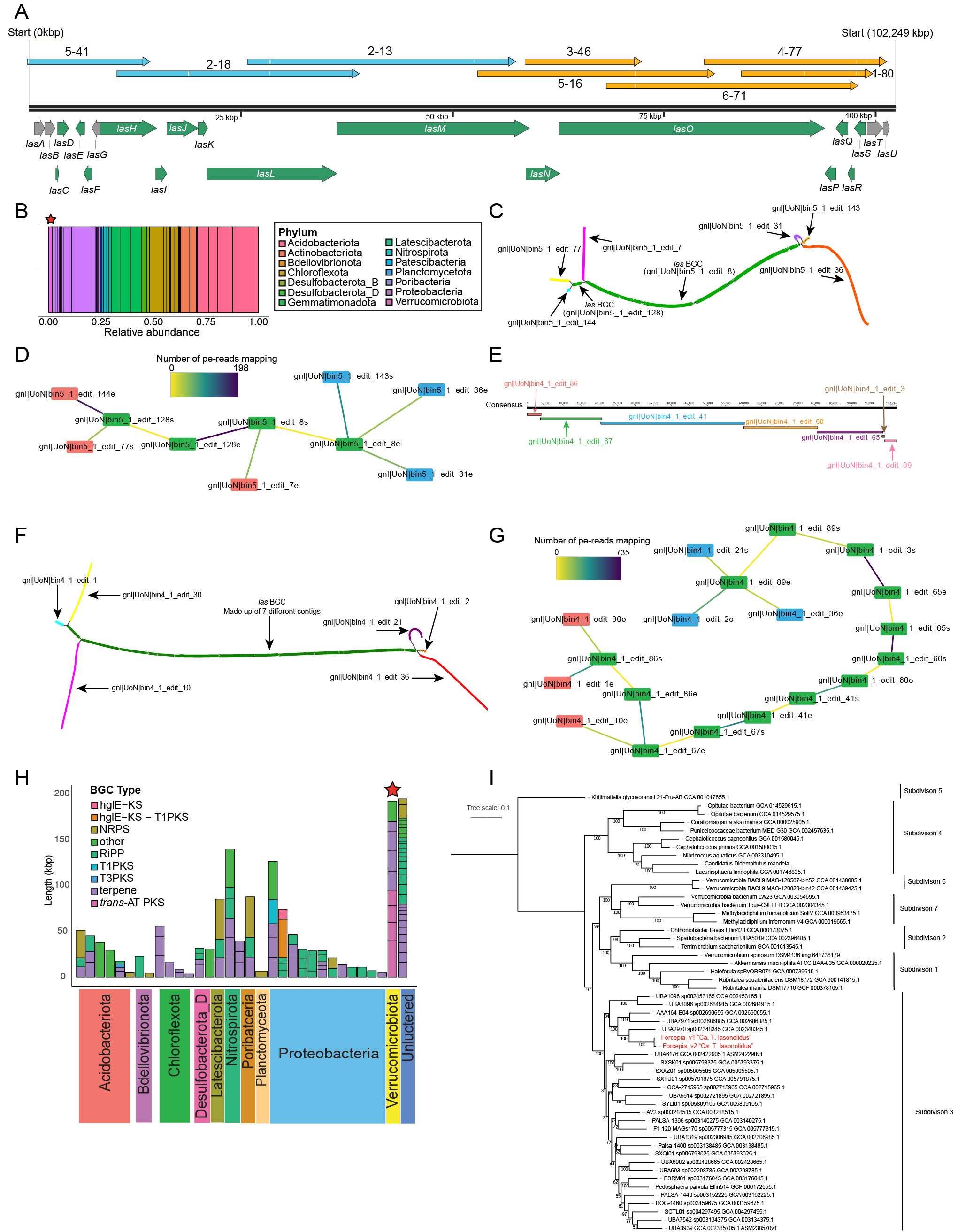

### Figure S2

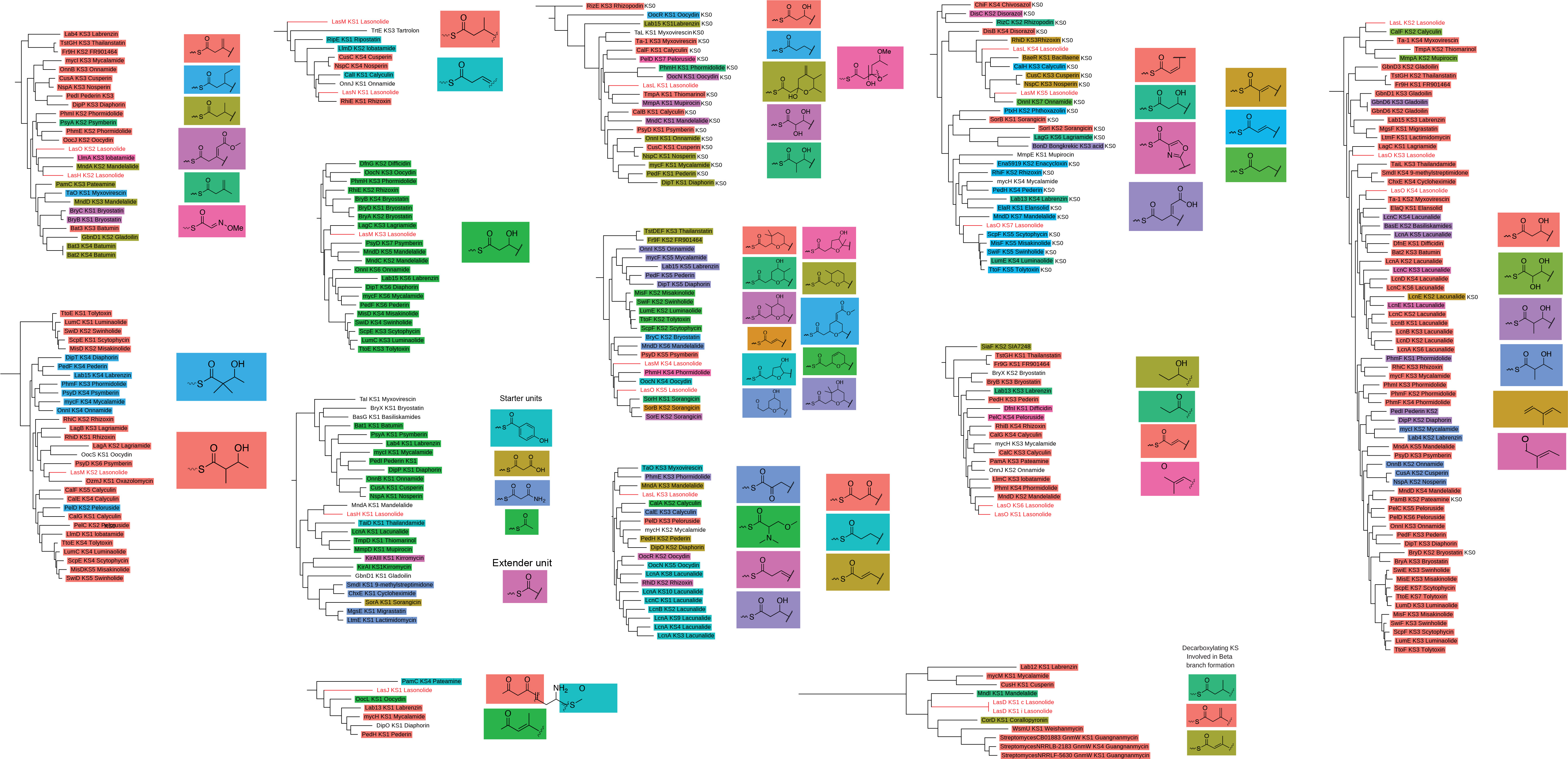

### Figure S3

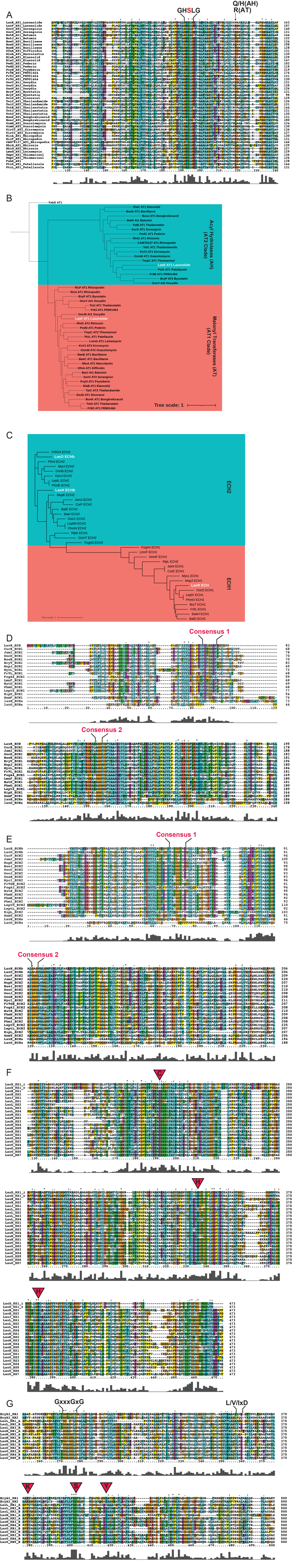

### Figure S4

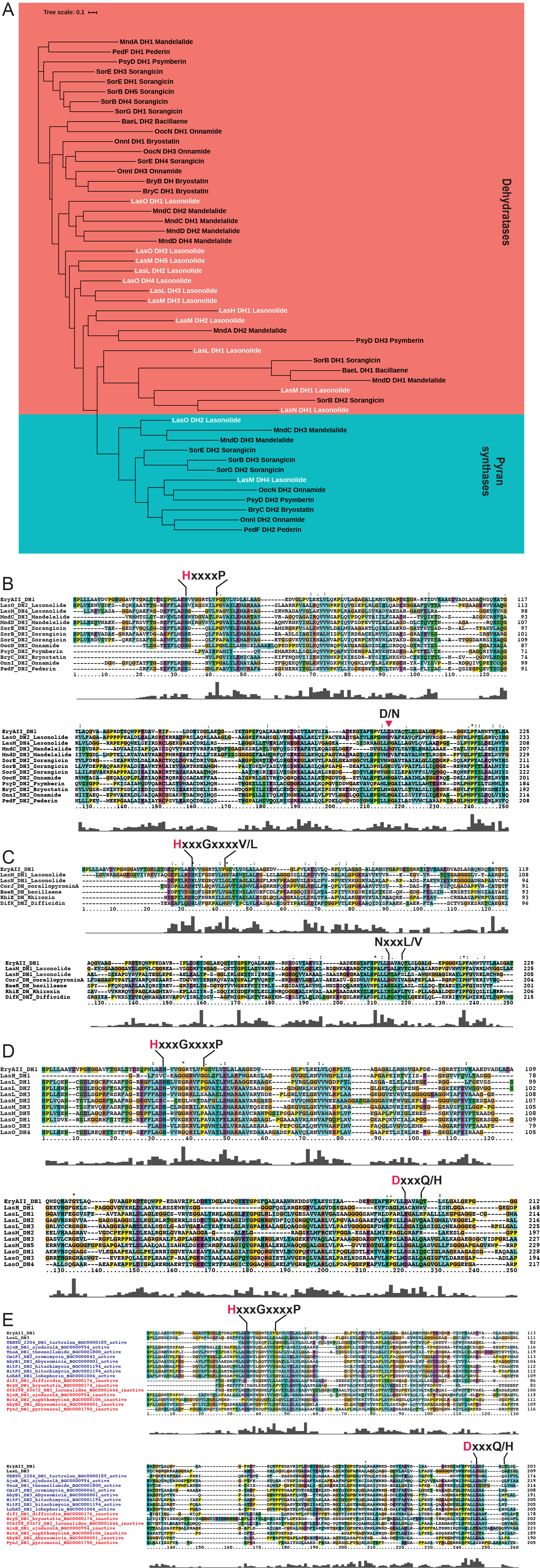

### Figure S5

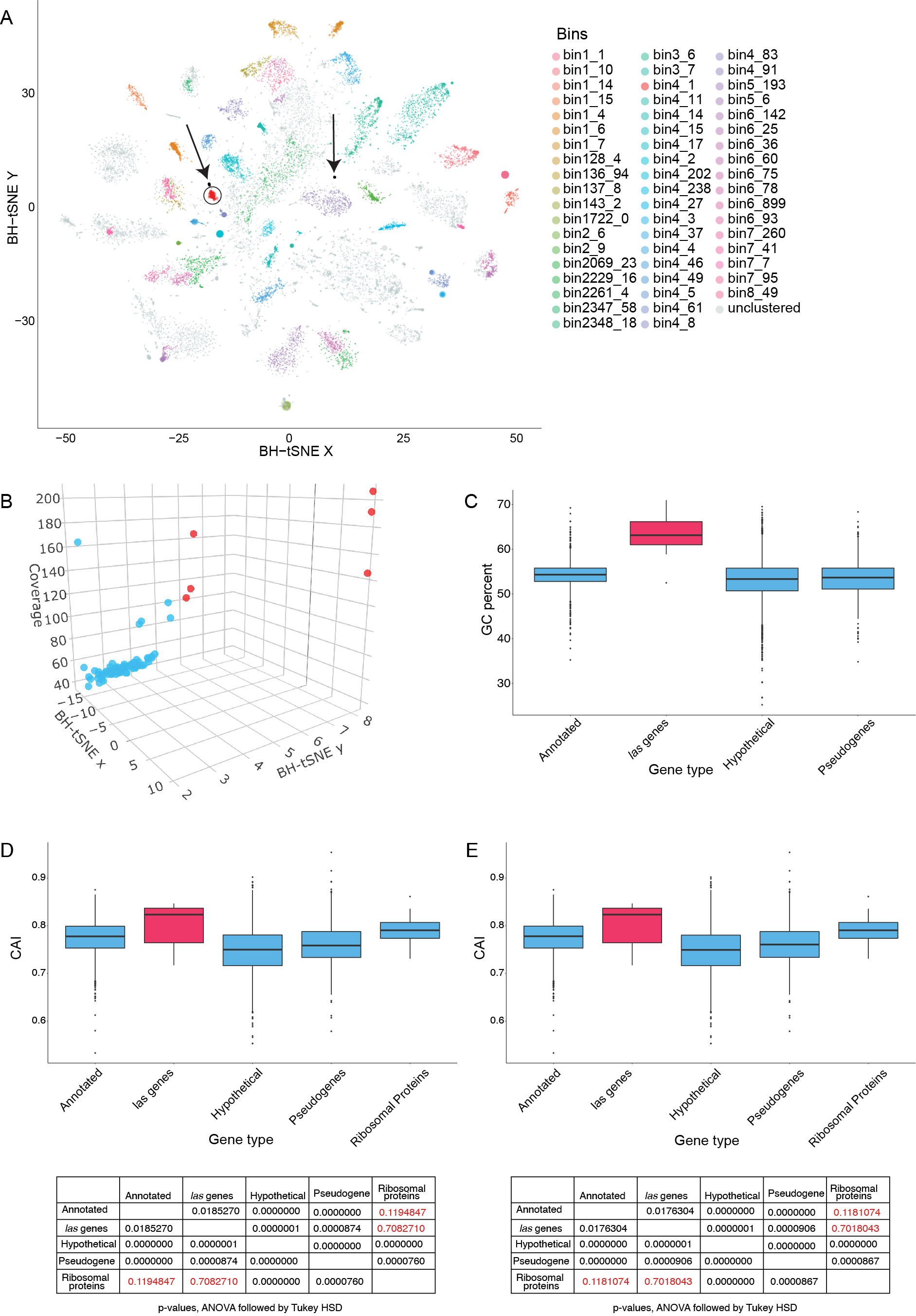
